# Exposure to the oral host niche yields rapid phenotypic and genotypic diversification in *Candida albicans*

**DOI:** 10.1101/254201

**Authors:** A. Forche, G. Cromie, A.C. Gerstein, N. V. Solis, Z. Pisithkul, W. Srifa, E. Jeffery, S.G. Filler, A.M. Dudley, J. Berman

**Affiliations:** Department of Biology, Bowdoin College, Brunswick, ME USA; Pacific Northwest Research Institute, Seattle, WA, USA; Department of Microbiology & Immunology, University of Minnesota, Minneapolis MN USA; Division of Infectious Diseases, Los Angeles Biomedical Research Institute at Harbor-UCLA Medical Center, Torrance, CA USA; Department of Medicine, David Geffen School of Medicine at UCLA, Los Angeles, CA USA; School of Molecular Cell Biology & Biotechnology, George S. Wise Faculty of Life Sciences, Tel Aviv University, Tel Aviv, Israel

**Keywords:** *Candida albicans*, aneuploidy, loss of heterozygosity, oropharyngeal Candidiasis, hypervariability, colony phenotype

## Abstract

*In vitro* studies suggest that stress may generate random standing variation, and that different cellular and ploidy states may evolve more rapidly under stress. Yet this idea has not been tested with pathogenic fungi growing within their host niche *in vivo*. Here, we analyzed the generation of both genotypic and phenotypic diversity during exposure of *Candida albicans* to the mouse oral cavity. Ploidy, aneuploidy, loss of heterozygosity (LOH) and recombination were determined using flow cytometry and ddRADseq. Colony phenotypic changes (CPs) in size and filamentous growth were evident without selection, and were enriched among colonies selected for LOH of the *GAL1* marker. Aneuploidy and LOH occurred on all chromosomes (Chrs), with aneuploidy more frequent for smaller Chrs and whole Chr LOH more frequent for larger Chrs. Large genome shifts in ploidy to haploidy often maintained one or more heterozygous disomic Chrs, consistent with random Chr missegregation events. Most isolates displayed several different types of genomic changes, suggesting that the oral environment rapidly generates diversity de novo. In sharp contrast, following *in vitro* propagation isolates were not enriched for multiple LOH events, except in those that underwent haploidization and/or had high levels of Chr loss. The frequency of events was overall 100 times higher for *C. albicans* populations following *in vivo* passage compared to *in vitro*. These hyperdiverse in vivo isolates likely provide *C. albicans* with the ability to adapt rapidly to the diversity of stress environments it encounters inside the host.

**Author summary:** Adaption is a continuous dynamic process that requires genotypic and phenotypic variation. Here we studied the effects of a single passage in a mouse oropharyngeal model of infection on the appearance of diversity in *C. albicans,* a common commensal of the human oral cavity and GI tract. We found that variation could be rapidly detected following oral colonization, with the frequency of genome change being considerably higher with pre-selection for recombination and colony phenotypic changes. Importantly, one third of all isolates had multiple genome changes, significantly higher than expected by chance alone. We suggest that some cells in the population are naturally hypervariable and that they are a major source of diversity upon which selection can act in stressful conditions *in vivo* and *in vitro*.

## Introduction

*Candida albicans* is a common commensal of the human GI tract and the oral cavity in healthy individuals, and also an opportunistic pathogen, especially in immunocompromised patients (CALDERONE 2012). In healthy people, the fungus is prevented from causing disease by the resident microbiota and the host immune system (LORENZ *et al.* 2004; RICHARDSON AND RAUTEMAA 2009). However, immune deficiencies or a minor imbalance of the microbiota (e.g., through administration of antibiotics) can be sufficient to cause superficial infections. During the course of infection, *C. albicans* encounters many different host environments to which it must adapt rapidly. Furthermore, it must cope with environmental fluctuations in established niches during long-term persistence in the host (STAIB *et al.* 2001). Determining the genetic and phenotypic changes that accompany the establishment of commensalism and the transition to pathogenicity (and hence how they can be prevented) is not known (NAGLIK *et al.* 2003; WILSON *et al.* 2009).

Several studies strongly suggest that *C. albicans* may have a very different arsenal of adaptation mechanisms when in direct contact with the host compared to laboratory conditions. For example, a novel cell phenotype (GUT) is unique to the commensal environment of the gastrointestinal tract (PANDE *et al.* 2013) and several genes (e.g., Cph2, Tec1) are specifically expressed under commensal conditions (ROSENBACH *et al.* 2010). Furthermore, cells in the commensal state express genes that suggest the presence of at least two sub-populations of exponentially growing cells alongside stationary-phase cells. In addition, the expression patterns of several genes are clearly distinct during growth *in vivo* vs *in vitro* (NOBILE *et al.* 2006; VANDEPUTTE *et al.* 2011; FANNING *et al.* 2012; LOHBERGER *et al.* 2014).

Recent studies in fungi found that genome instability caused by large-scale chromosomal changes, including gross chromosomal rearrangements (GCRs), supernumerary Chrs (SNCs), aneuploidy and loss of heterozygosity (LOH), are more frequent under stress conditions *in vitro* and *in vivo* (RUSTCHENKO *et al.* 1997; SELMECKI *et al.* 2006; COYLE AND KROLL 2008; POLAKOVA *et al.* 2009; FORCHE *et al.* 2009a; FORCHE *et al.* 2011; HICKMAN *et al.* 2013). Aneuploidy in particular has been shown to be one of the mechanisms that can lead to antifungal drug resistance in pathogenic fungi including *Crypococcus neoformans* and *Candida glabrata* (SELMECKI *et al.* 2006; POLAKOVA *et al.* 2009; SIONOV *et al.* 2010). Interestingly, a recent study showed that *C. albicans* forms very large cells in response to acute micronutrient limitation, in particular to zinc. Cell size has been shown to be correlated with ploidy (HICKMAN *et al.* 2013) and indeed flow cytometry data support that these gigantic cells may be aneuploid (MALAVIA *et al.* 2017). In *C. neoformans,* Titan cells are large polyploid cells that can rapidly produce drug resistant aneuploid daughters upon exposure to the drug fluconazole (GERSTEIN *et al.* 2015), supporting the idea that aneuploidy is a common adaptation mechanism of pathogenic fungi.

Our previous study of ∼80 *C. albicans* isolates recovered from the mouse model of hematogenously disseminated candidiasis (BSI) from mice kidneys (FORCHE *et al.* 2003; FORCHE *et al.* 2009a) provided a first glimpse into the types of genomic changes that *C. albicans* undergoes at the population level. We discovered higher rates of phenotypic and Chr-level genetic variation following passage of *C. albicans in vivo* relative to passage *in vitro*. In addition, missegregation events, including whole Chr aneuploidy and LOH, were positively associated with altered CPs.

The oral cavity is one of the few host niches that is both a commensal and pathogenic niche (VARGAS AND JOLY 2002; PATIL *et al.* 2015). *C. albicans* has been found as part of the commensal microflora in up to two thirds of the healthy population (VILLAR AND DONGARI-BAGTZOGLOU 2008; PANKHURST 2013). Oral and oropharyngeal candidiasis can develop as consequence of developed immunodeficiency (e.g. HIV/AIDS), underlying diseases such as diabetes, and treatment with broad-spectrum antibiotic, corticosteroids and chemotherapy (SOBUE *et al.*; LYON *et al.* 2006; LU *et al.* 2017). In the oral niche, fungal-host interactions are highly dynamic due to a multitude of factors including the presence of antimicrobial salivary peptides and the microbiota of bacterial and fungal species that co-exist and compete for nutrients on epithelial cells (DEMUYSER *et al.* 2014; JAKUBOVICS 2015) and the highly fluctuating environmental conditions (e.g., temperature, pH) (PARK *et al.* 2009). Unexpectedly, we recently identified haploid, mating-competent *C. albicans* isolates for the first time, and most of these haploids were recovered after *in vivo* passage in an oral model of infection (HICKMAN *et al.* 2013). This extraordinary finding highlights the contribution of and the need for *in vivo* studies to the discovery of novel aspects of *Candida* biology in general and of host-pathogen interactions in particular.

To further our understanding on the acquisition of standing variation of *C. albicans* during infection, we performed experimental evolution of *Candida albicans* in mice to analyze the appearance of genotypic and phenotypic diversity during passage through the mouse oral cavity for 1, 2, 3 or 5 days using an oropharyngeal model of infection (KAMAI *et al.* 2001; SOLIS AND FILLER 2012). We found that diversity is rapidly generated after exposure to the oral host niche, and that many of these changes are identified in multiple mice. The overall high within-mouse diversity and multiple changes per isolate was high independent of the duration of infection. Surprisingly, the generation of multiple genetic changes in a single isolate appears to occur with higher frequency than would be expected by random chance alone. Taken together, our results suggest that exposure to the host (and/or the transition from *in vitro* to *in vivo* growth conditions) generates highly variable isolates at a frequency 2 orders of magnitude higher than *in vitro*.

## Methods

### Isolate maintenance and DNA extraction

Strains used to generate parental strain YJB9318 are listed on Table S1. YJB9318 and recovered isolates were grown on YPD (2% glucose, 1% yeast extract, 1% bacto peptone, 20 mg/L uridine with 1.5% agar added for plate cultures). Gal phenotypes were assessed on MIN-Gal (0.67% yeast nitrogen base without amino acids, 2% galactose, 1.5% agar; only Gal^+^ isolates grow) and 2-deoxygalactose medium (2DOG; 0.1% 2-deoxygalactose, 0.5% raffinose, glycerol, 1.5% agar; only Gal^-^ isolates grow). All isolates are stored long-term in 50% glycerol at -80°C. DNA extractions were performed as described previously (SELMECKI *et al.* 2005).

### Construction of strain YJB9318

Plasmids and primers used in this study are listed in Table S1. YJB9318 is a derivative of strain RM1000 #2 (Table S1) in which one copy of *GAL1* was replaced with *URA3* (*GAL1/Δgal1*::*URA3)*. First, the *URA3* marker was amplified from plasmid p1374 with primers 1672 and 1673 (Table S1), and transformed into isolate YJB7617 (RM1000#2) replacing one copy of *GAL1* (YJB8742) (LEGRAND *et al.* 2008). Correct disruption of *GAL1* was confirmed by diagnostic PCR using primers 1674 and 1675 (Table S1). To make YJB9318 prototrophic, *HIS1* was reintroduced into strain YJB8742 by transforming with plasmid p1375 (pGEM-*HIS1*) that was cut with restriction enzyme *Nru*I. Diagnostic PCR with primers 728 and 565 (Table S1) confirmed correct integration of *HIS1* at its native locus. To ensure that transformation did not cause any genomic changes to the parental strain, single nucleotide polymorphism (SNP) microarrays and SNP/Comparative genome hybridization arrays (SNP/CGH) were performed as described previously (data not shown) (SELMECKI *et al.* 2005; FORCHE *et al.* 2009a; ABBEY *et al.* 2011).

### PCR conditions for transformation and diagnostic PCR

PCRs for transformation were performed in a total volume of 50 μl with10 mM Tris–HCl (pH 8.0), 50 mM KCl, 1.5 mM MgCl_2_, 200 μM each dATP, dCTP, dGTP, and dTTP, 2.5 U rTaq polymerase (TAKARA), 4 μl of 10 μM stock solution of each primer, and 1.0 μl of template (p1374). PCRs were carried out for 34 cycles as followed: initial denaturation step for 5 min at 94°C, denaturation step for 1 min at 94°C, primer annealing step for 30 s at 55°C, extension step for 1 min at 72°C, and a final extension step for 10 min at 72°C. Each PCR product was checked by gel electrophoresis for the amplification of the desired PCR fragment. PCR products were purified using ethanol precipitation.

Diagnostic PCR was performed in a final volume of 25 μl with 10 mM Tris–HCl (pH 8.0), 50 mM KCl, 1.5 mM MgCl_2_, 100 μM each dATP,dCTP, dGTP, and dTTP, 2.5 U rTaq polymerase, 2 μl of 10 μM stock solution of each primer, and 2.5 μl genomic DNA. PCRs were carried out for 30 cycles as followed: initial denaturation for 3 min at 94°C, denaturation step or 1 min at 94°C, primer annealing step for 30 s at 55°C, extension step for 1 min at 72°C, and a final extension step for 5 min at 72°C. Five microliters of PCR product was run on a 1% agarose gel to verify that the fragment was of the appropriate size.

### Model of oropharyngeal Candidiasis (OPC)

The OPC model was essentially performed as described previously (SOLIS AND FILLER 2012). Briefly, male BALB/c mice (21-25 g; Taconic Farms) were immune-suppressed with cortisone acetate (225 mg/kg, Sigma) on days -1, 1, and 3 of infection. For inoculum preparation, strain YJB9318 was grown in MIN-Gal medium to ensure that no Gal^-^ cells arose prior infection. A total of twenty mice were infected with 1 × 10^6^ cells of strain YJB9318 (Table1). Of these, 17 mice survived to the scheduled dates of sacrifice. On days 1, 2, 3, and 5 post-infection (FIG.1, FIG.S1), 4-5 mice were euthanized. The tongues were extracted, weighted, and homogenized. Next, appropriate dilutions were spread onto YPD agar plates for total CFU counts and onto 2DOG agar plates to determine the number of Gal^-^ cells. Recovered isolates were directly picked from the original YPD and 2DOG plates to 96well plates with 50% glycerol and stored at -80°C to avoid any changes to the isolates not acquired during *in vivo* passage.

**FIG.1.**
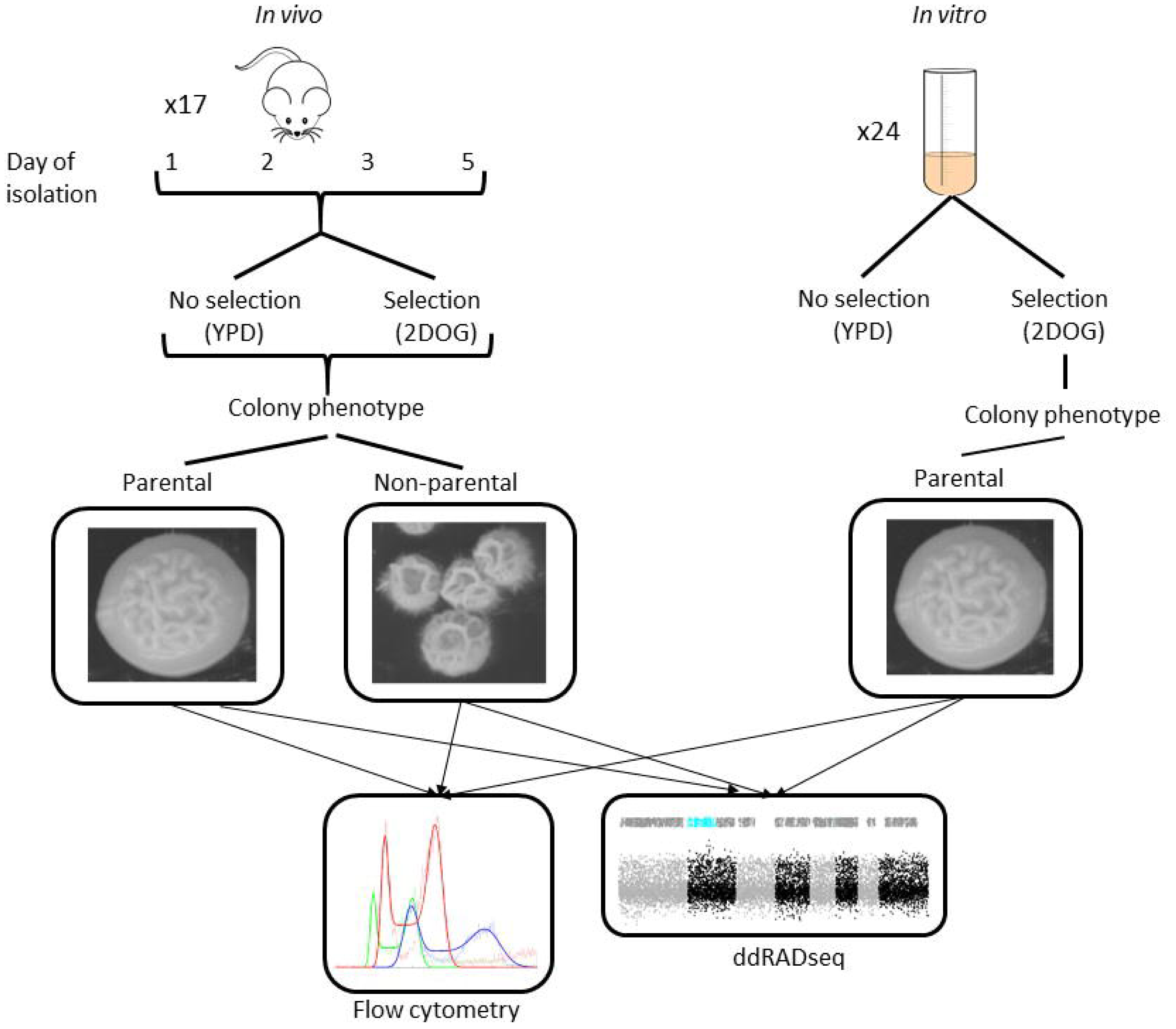
Experimental overview of *in vivo* and *in vitro* experiments.

**Table 1.** Frequency of colony morphology phenotypes.

| CM* | Total<br>N = 429 | Day1<br>N = 25 | Day2<br>N = 54 | Day3<br>N = 195 | Day5<br>N = 155 |
| --- | --- | --- | --- | --- | --- |
| 00 | 0.62 | 0.8 | 0.74 | 0.66 | 0.49 |
| 01 | 0.03 | 0 | 0.04 | 0.04 | 0.02 |
| 02 | 0.03 | 0.04 | 0.04 | 0.02 | 0.05 |
| 03 | 0.01 | 0 | 0.00 | 0.02 | 0.01 |
| 10 | 0.06 | 0.04 | 0.07 | 0.04 | 0.10 |
| 11 | 0.07 | 0.04 | 0.02 | 0.05 | 0.12 |
| 12 | 0.14 | 0.08 | 0.06 | 0.12 | 0.21 |
| 13 | 0.01 | 0 | 0 | 0.02 | 0.01 |
| 20 | 0.02 | 0 | 0.04 | 0.04 | 0 |

\*Colony morphology, see Fig. 2D for representative images.

To confirm the Gal status of recovered isolates, they were grown overnight in deep 96-well plates containing 300 μl YPD broth. Cultures were washed once with distilled water, 5 μl of each culture were spotted onto 150 x 100 mm YPD plates, MIN-Gal and 2DOG medium, and plates were incubated for 2 days at 30^°^C to assess growth. The frequency of LOH at the *GAL1* locus was determined using the ratio of total isolates recovered (CFUs on YPD) divided by the total number of 2DOG^R^ isolates. The *in vitro* frequency of *GAL1* loss in strain YJB9318 was measured as described previously (FORCHE *et al.* 2009).

### Assessment of colony phenotypes (CPs) and selection of isolates for genotypic analysis

Previously, we showed that isolates with missegregation events (whole Chr aneuploidy and whole Chr LOH) exhibited CPs consistent with slow growth and abnormal filamentous growth (FORCHE *et al.* 2005; FORCHE *et al.* 2009). To increase the ability to identify genotypic changes, we plated all isolates for CPs on YPD at 30°C and scored single colonies after 3 days. CPs were determined for colony diameter (smaller or larger than parental strain, first number) and filamentous growth (degree of wrinkling compared to parental strain, second number) resulting in a binary code for each of the 7 unique CPs (see FIG.2D for representative images). For further genotypic analysis, all isolates with altered CPs (6 Gal^+^ and 158 Gal^-^), and isolates with parental CP (148 Gal^+^ and 116 Gal^-^) from a total of 17 mice were chosen to yield a set of 429 isolates (FIG.S1, Table S2).

**FIG.2.**
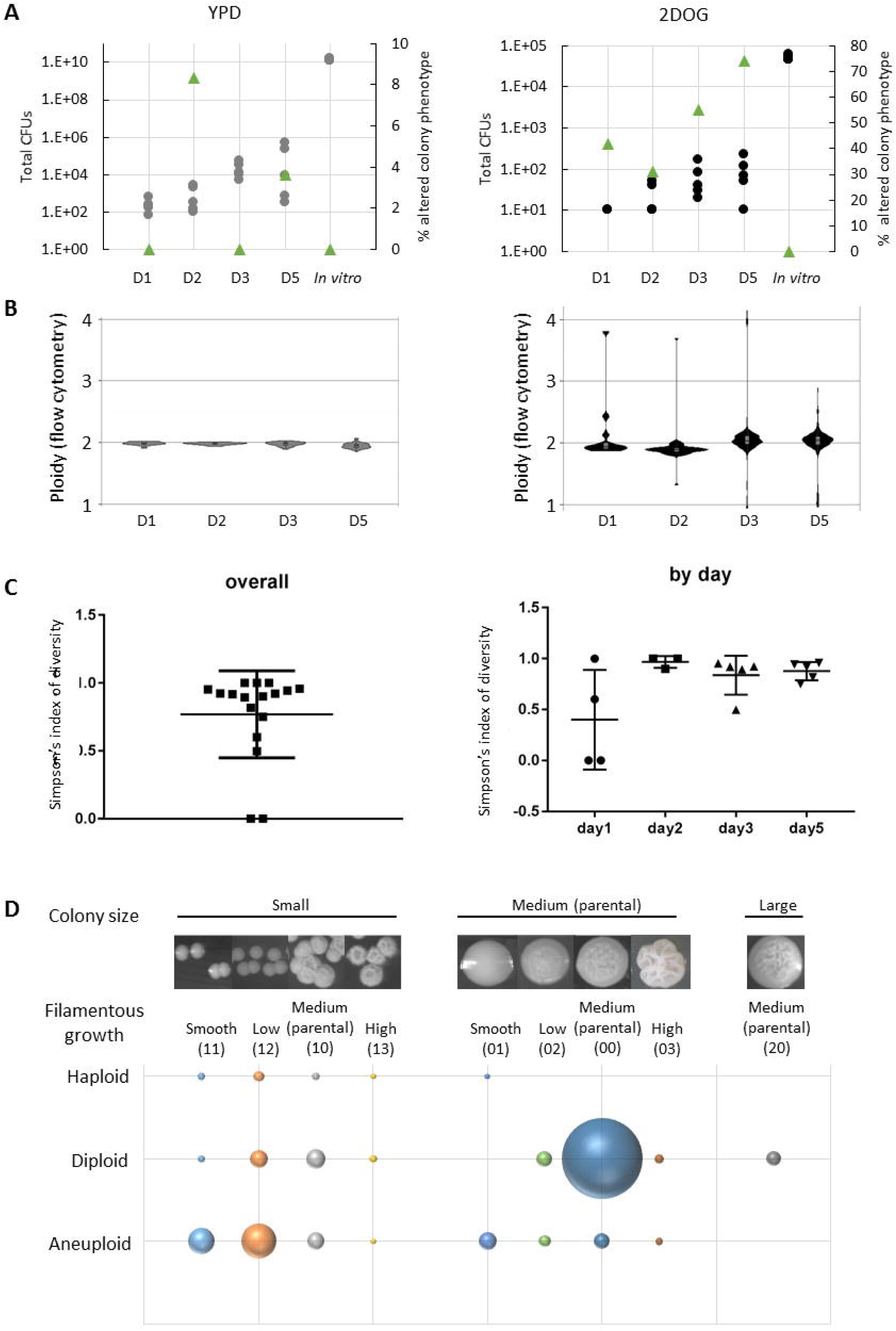
Genotypic and phenotypic diversity arises early during oral infection. **A.** CPs arise later in the Gal^+^ group (day 2, left graph) compared to the Gal^-^ group (day 1, right graph) but are not observed *in vitro*. **B.** Ploidy changes (as measured by flow cytometry) do not arise in the Gal^+^ group, but do so in the Gal^-^ group with ploidy shifts observed as early as day 1 post infection. **C.** Simpson’s D is high on average and depends strongly on composition of populations within mice (Gal^+^/Gal^-^ ration of isolates) left, overall Simpson’s D; right, Simpson’s D by days spent in mice **D.** Bubble plot shows the number of haploid, diploid or aneuploid isolates (as measure by flow cytometry) that exhibit indicated CPs determined by growth on YPD at 30°C for 3 days. CP binary codes are shown in parenthesis (see also Table S2). Bubble size reflects the number of isolates. Circles, total CFUs, green triangles, CPs.

### Determination of ploidy by flow cytometry

Ploidy of all recovered isolates was determined as described previously (ABBEY *et al.* 2011). Briefly, each isolate was streaked out to single colony onto YPD plates and incubated for 3 days at 30°C. Single colonies were transferred to deep 96-well plates containing 0.6 ml of YPD and cultures were grown overnight (16 hrs) at 300 rpm to stationary phase. Fifty microliters were transferred to new deep 96-well plates containing 250 μl YPD broth, and cultures were grown for 6 hrs at 30°C at 300 rpm. Two hundred fifty microliters of culture was transferred to round bottom 96-well plates, cells were spun down at 1,000 rpm, and resuspended in 20 μl of 50:50 buffer (50 mM Tris HCl, pH 8.0, 50 mM EDTA, pH 8.0). To fix cells, 180 μl of 95% ethanol was added to each well. Cells were treated with 0.1 μg/ml RNase (1 hr at 37°C) and 5 mg/ml Proteinase K (30 min at 37°C) followed by staining with SybrGreen for 1hr in the dark. After a final wash in 50:50 buffer, cells were resuspended in 50:50 buffer and run on a flow cytometer (FACSCALIBUR). A customized MATLAB script was used to calculate ploidy for each isolate using a diploid and a tetraploid isolate as controls (ABBEY *et al.* 2011).

### Whole genome karyotyping using double digest restriction site associated DNA sequencing (ddRADseq)

ddRADseq was carried out using the restriction enzymes *Mfe*I and *Mbo*I (LUDLOW *et al.* 2013). For each lane of Illumina sequencing (up to 576 isolates/lane), raw read sequences were split into isolate-specific pools based on their associated 6 bp TruSeq multiplex and 4 bp inline barcode sequences, allowing 1 mismatch in the i7 barcodes and no mismatches in the inline barcodes. A minimum barcode quality of Phred = 20 was applied to all bases of the inline barcode. Reads were then aligned to the *C. albicans* reference (SC5314 v. A21-s02-m04-r01) using BWA (v.0.7.5) allowing 6 mismatches and quality trimming using the parameter -q 20. The SAMtools (v.0.1.17) (LI *et al.* 2009) mpileup command was then used to create a pileup file for each isolate, using the -q 20 and –C 50 parameters. From the pileup file, the count of all observed bases at each covered reference position was calculated.

#### Ploidy Estimation from ddRADseq

From the aligned-read SAM file, the position of the *Mfe*I end of each read was determined (5’ end of forward reads, 3’ end of reverse reads). Only reads with a Phred-scaled mapping alignment quality of at least 20 were considered. The occurrence of each end position was then counted, resulting in a set of marker positions for each isolate along with the number of reads aligning to each of those positions. For all isolates in a sequencing run, a matrix of observed read counts at all positions was generated. Because of the long tail of infrequently observed sites, counts of 1 were treated as counts of zero. Only positions counted in at least one isolate were retained. The matrix of counts was then edited to remove any isolates with ≥ 20% positions with 0 counts. Following this, marker positions were filtered to only include those occurring in >80% of remaining isolates. The edited count matrix was used to calculate relative ploidy at each marker position as follows. Each isolate was normalized for depth of sequencing by dividing all observed counts by the median value of all counts > 0. To control for marker-to-marker variation in coverage (largely due to the size of the associated DNA fragment), normalized coverage at each marker in each isolate was divided by the median coverage (ignoring zero values) of that marker across a set of control euploid isolates. Before plotting, globally noisy markers, with standard deviations >0.5 across all isolates (not including zero values), were removed and for each isolate a minimum raw count coverage of 10 or 20 was required at each position to remove low-confidence estimates for that isolate.

Ploidy was also assessed on a by-Chr basis. The unedited, original counts matrix was used to calculate the proportion of all reads in each isolate aligning to each Chr. The value for each Chr was then normalized by dividing the median value for that Chr across the set of euploid control isolates. To account for variation in genome size in aneuploids, values for each isolate were then further normalized by dividing each Chr value by the median Chr value for that isolate. For euploid isolates this should produce values of ∼1 across all Chr. A diploid with one trisomic Chr would have a value of ∼1.5 for one Chr and 1 for the rest. On the assumption that most isolates are essentially diploid, values were converted into ploidy by multiplying by 2, 3 or 4 in the small number of isolates where this produced Chr copy number values closer to whole numbers.

#### Estimation of Allele Ratios

Heterozygosity was assessed at all heterozygous sites in parental strain SC5314 (MUZZEY *et al.* 2013) ignoring indels. For each Chr, SC5314 heterozygous sites were identified by aligning the two phased haploytpes (“A” and “B”) from (MUZZEY *et al.* 2013) using the Mummer (v3.22) nucmer command with the parameters ‐c 100 ‐l 10 ‐b 200. Alignments were filtered using the delta-filter command with the –g parameter and snps were then called using the show-snps command with the parameters ‐r ‐C ‐H –T. the counts of the two expected alleles were extracted from the count of all observed alleles (as above) in each isolate for every expected het site covered by reads. The binomial probability of the observed counts was then calculated using models 1A:0B (homozygous A), 3A:1B, 2A:1B, 1A:1B, 1A:2B, 1A:3B and 0A:1B (homozygous B). For example for model 3A:1B the binomial probability of the observed data would be calculated based on P(A) = 0.75, P(B) = 0.25. For the homozygous models, observed counts of the “wrong” allele were assumed to be errors with a probability of 0.01, with the expected allele having a probability of 0.99. For each site, the set of possible models was then consolidated based on the copy number of the Chr, identified from the whole Chr read proportion analysis above. For disomic Chrs, models 1A:0B, 1A:1B and 0A:1B were compared and the best and second best models identified. For trisomic Chrs, the best and second best models were identified from models 1A:0B, 2A:1B, 1A:2B and 0A:1B. For tetrasomes, the models compared were 1A:0B, 3A:1B, 1A:1B, 1A:3B and 0A:1B. After identification of the best model, for each site in each isolate a LOD score (Log10 P (Best Model)/P(Second Best Model) was then calculated. For each dataset, markers were removed unless they were classified as heterozygous (best model = 1A:1B with LOD > 1) in at least one isolate. For visualization, a median sliding window of size 7 was applied to the best model values, ordered by genome position.

### Population frequency of changes

To obtain estimates of the population frequency of CPs and genotypic changes (i.e., not just for the analyzed isolates), we calculated the frequencies of phenotypic and genotypic changes for Gal+ and Gal‐ isolates based on the total number of *C. albicans* CFUs from the experiment (8.52 x 10^5^), the number of Gal^+^ isolates (8.51 x 10^5^; 99.89% of the total), and the number of Gal^-^ cells (9.1 x 10^2^; 0.11%). Extrapolation showed that the frequency of Gal^+^+ CP in the total population was 3.9 x 10^-2^ (∼1/400 cells) and ∼5.8 x 10^-4^ for Gal^-^+CP (∼6/10,000 cells) (FIG.S1). The frequency of genotypic changes (Chr1 changes excluded) for Gal^+^ cells was 13 x 10^-2^ and 5.2 x 10^-4^ for Gal^-^ cells. (FIG.S1).

A customized MATLAB script was written to calculate the total likelihood for all the permutations for each number of genotypic changes. A chi square goodness-of-fit test was applied to ask whether the difference between the expected and observed numbers was due to sampling variation, or whether it was a real difference. A p-value of significant.

### Diversity index

ddRADseq data was used to determine the number of unique karyotypes that were present in each mouse and the number of colonies that exhibited each karyotype. A karyotype was considered unique if there was either a unique whole or partial Chr aneuploidy or LOH event and/or there was variation in the LOH breakpoint, compared to other colonies isolated from the same mouse. Diversity was then calculated as Simpsons index 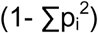 (SIMPSON 1949), where p_i_ is the proportional abundance of each colony type (note that if there is only one karyotype, diversity is 0).

## Results

To analyze diversification rates of *C. albicans* on a mucosal surface, we seeded the oral cavity of 20 corticosteroid-treated mice with 10^6^ cells originating from YJB9318, a single, colony-purified *C. albicans* strain that was heterozygous for *GAL1* (GORMAN *et al.* 1992). Groups of 4-5 mice were sacrificed on days 1, 2, 3 and 5 and isolates were recovered from tongue homogenates (FIG.1 and FIG.S1). Initiating the experiment with a Gal+/- strain enabled us to acquire evolved isolates from both YPD plates (Gal^+^, 541 colony isolates, no selection, unknown genomic changes) and 2-deoxygalactose (2DOG) plates (Gal^-^, 360 colony isolates, minimum genomic change of LOH at the *GAL1* locus) (FIG. 2A and S1). All recovered isolates were first screened for changes in colony phenotype (CPs) and we identified 7 distinct CPs, 3 of them were also detected among Gal^+^ isolates and all 7 were detected among Gal^-^ isolates (FIG.S1, Fig.2, and see below). As measured by flow cytometry, the majority (72%) of isolates retained a diploid genome content (FIG. 2B). Strikingly, eleven isolates had haploid or near-haploid genome (2%) content. An additional seven isolates were tetraploid or near-tetraploid while the remainder (26%) had ploidy values consistent with aneuploid diploids (Table S2).

Twenty-four hours after infection, the oral fungal burden was approximately 10^2^ CFUs per g tissue, suggesting that only a small proportion of the starting inoculum initiated the oral infections (FIG. 2A). The number of CFUs generally increased with time of infection and the proportion with CPs increased slightly (Table 1). The proportion of Gal1^-^ CFU increased proportional with the total CFU (measured by comparing the frequency of 2DOG^R^ isolates and YPD isolates). The overall frequency of LOH at *GAL1* was two orders of magnitude higher *in vivo* compared to *in vitro* (FIG.2A, S2). Ploidy changes were much more prevalent in the isolates selected for *GAL1* LOH (FIG. 2B) compared to isolates from YPD and non-diploid isolates were frequently associated with reductions in both colony size and filamentous growth (FIG. 2D).

## Genotypic diversity by RADSeq

ddRADseq analysis was used to analyze Chr copy number and allele frequencies from 154 isolates off YPD and 275 isolates off 2DOG. The overall diversity was calculated using Simpsons index of diversity (1-D) (see methods) for each mouse. The within-mouse diversity was variable at day 1 and remained high thereafter (FIG. 2C). This suggests that diversity was generated early after infection and was not highly deleterious to survival and growth within the host.

We then clone-corrected the data set based upon the assumption that isolates with identical genotype and CP from the same mouse were likely to be daughter isolates resulting from a single mutational event. When the same event (genomic change) was found in different mice, we expect that event was either frequent and/or subject to strong selective pressure in the mice and was termed a ‘recurrent’ event.

Isolates that underwent ploidy shifts based on flow cytometry were re-analyzed by ddRADseq. Interestingly, euploid shifts (the loss or gain of complete sets of Chrs) were extremely rare; only three (of 10 confirmed) haploids and none of the 7 confirmed triploids or tetraploids were euploid (FIG. 3A and B). By contrast, trisomy was detected for every Chr, with higher trisomic frequencies for smaller Chrs and ChrR (FIG.4A and B) (Table 2, S3). There were seven isolates in which the majority of Chrs (>4) were non-disomic or where Chrs were present in multiple ploidy levels (e.g. monosomy, disomy and trisomy within the same isolate), providing indirect evidence that euploid shifts likely preceded subsequent Chr missegregation events (FIG.3C) (FORCHE *et al.* 2008; HARRISON *et al.* 2014; HICKMAN *et al.* 2015). Importantly, aneuploidy was detected in both Gal^+^ and Gal^-^ isolates (FIG.S3).

**FIG.3.**
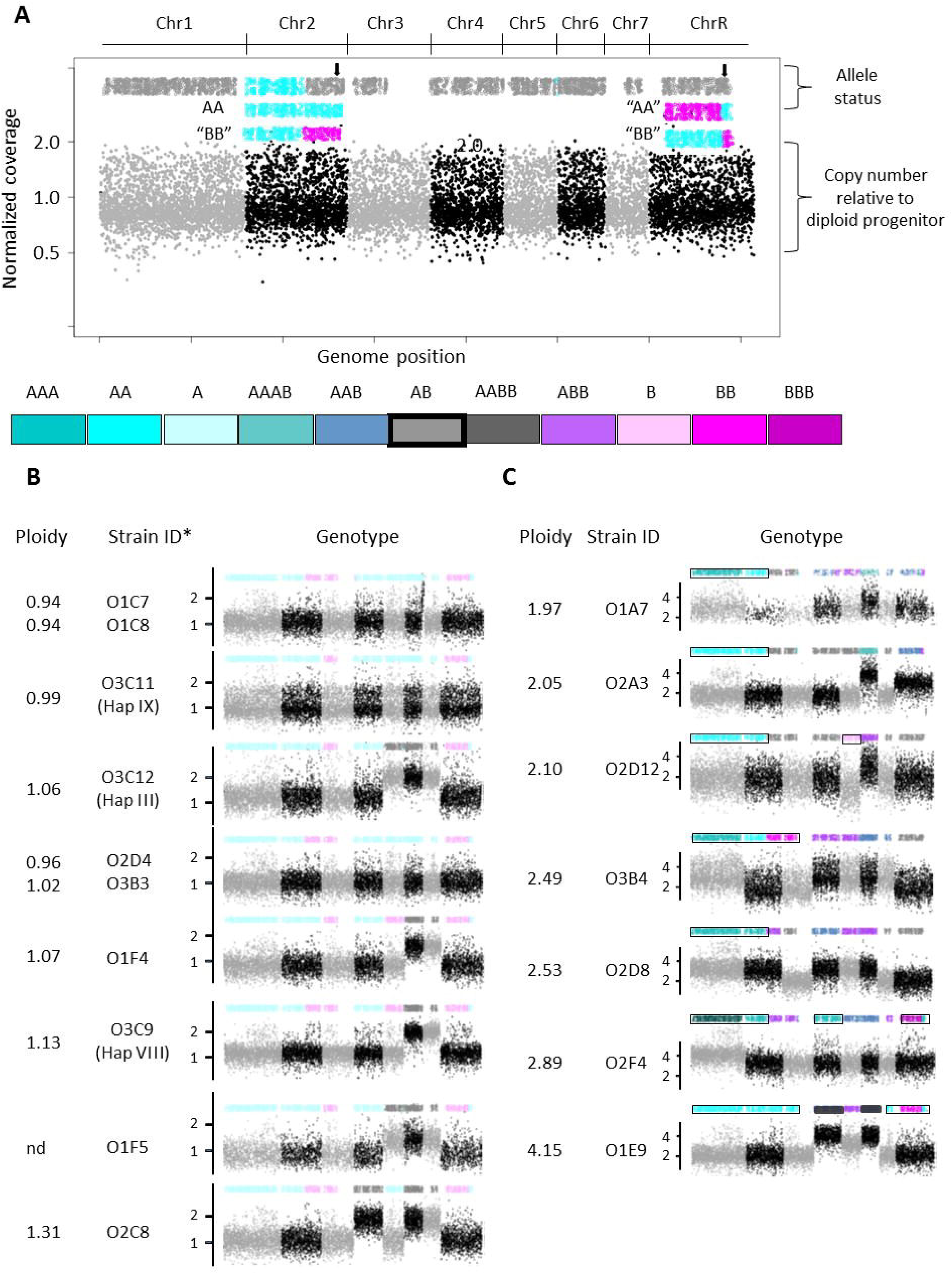
Whole genome ploidy shifts are rare.**A**. Shown is the ddRADseq whole genome karyotype for parental strain YJB9318. Allele status is indicated on the top. Bottom half of figure provides copy number for each Chr relative to diploid parent (1 = 2 copies) Chrs are colored in light grey and black to indicate start/end of each Chr. Color-coding is used throughout for indicated genotypes. Each dot on the lower part (copy number) is a copy number estimate for a restriction fragment based on the reads aligning to one end of the restriction fragment. The dots on the upper part (allele status) are maximum likelihood estimates of allele ratios at each (sequenced) known SNP site, constrained by the Chr or segment copy number and smoothed across x number of adjacent sites (see also methods). The colors for the allele status provide exact genotype for each Chr. Note: This strain background (RM1000 #2) has a preexisting Chr2L allele A homozygosis and a crossover on ChrR occurred during generation of the parental strain that was unmasked in isolates that became homozygous (see red arrow). In the case of whole Chr LOH, the genotype at the centromere was called (see black arrows). Gaps in allele coverage on Chrs3, 7, and R are due to lack of heterozygosity in the reference strain SC5314 used for analysis (FORCHE *et al.* 2004; VAN HET HOOG *et al.* 2007; BUTLER *et al.* 2009). **B.** Haploids and near haploids exhibit different genotypes, *(strain names in parenthesis from Hickman *et al.* 2013), y-axis, Chr copy number, x-axis, Chrs are ordered Chr1-7, and R. **C.** Isolates with > 2 ploidies/genome suggestive of ploidy shifts in progress.

**FIG.4.**
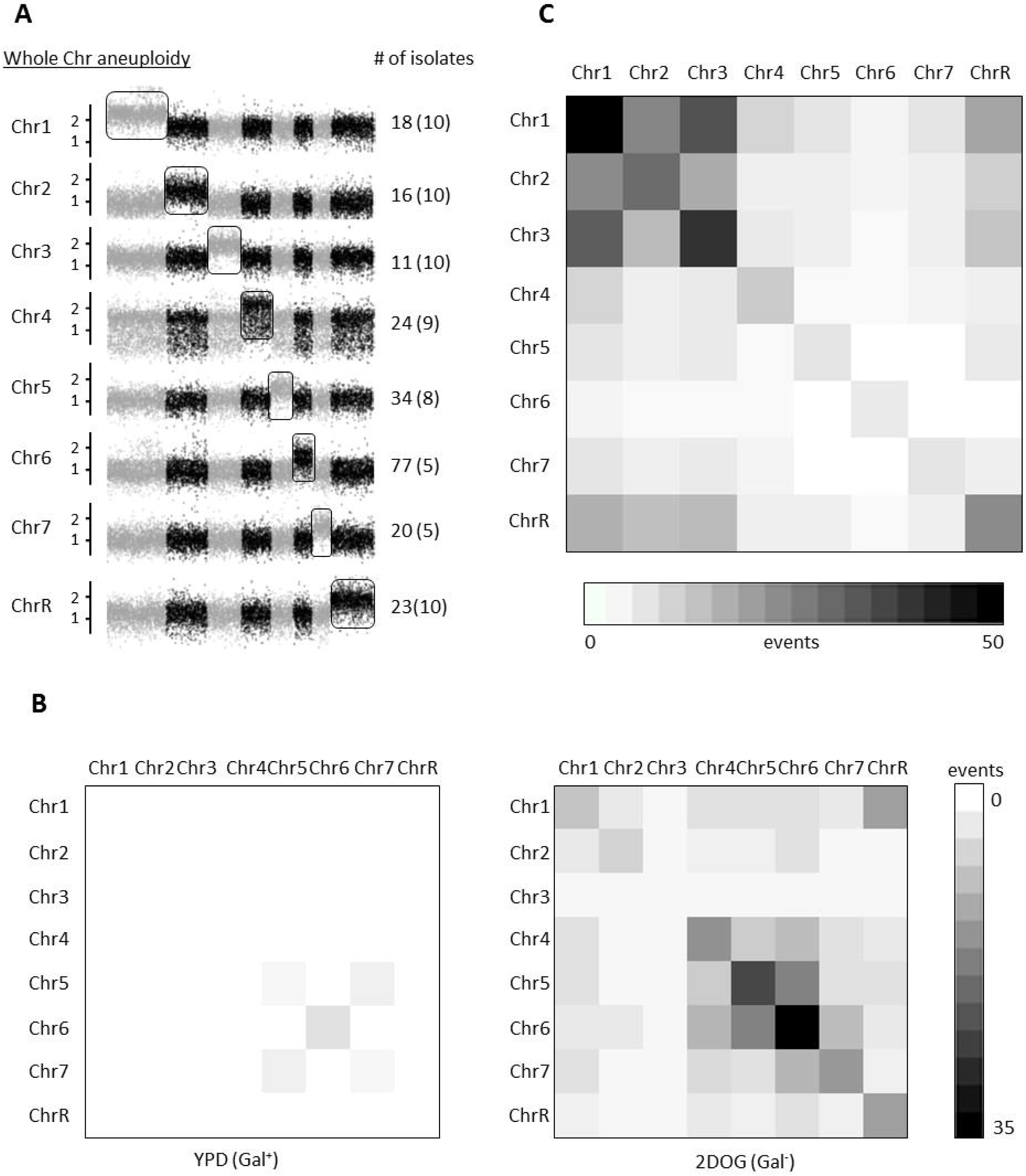
Overview of missegregation events across Chrs. **A.** Whole Chr aneuploidies include trisomies and tetrasomies. The number of disomic Chrs from haploids and near haploids is shown in parentheses. Y-axis: normalized copy number relative to diploid parent. Aneuploid Chrs are boxed in. Note: Images show single whole Chr aneuploidies for clarity; most isolates carry more than one whole Chr aneuploidy. **B.** Single and double aneuploidies are detected both for Gal^+^ and Gal^-^ isolates. Shown is number of isolates with 1 aneuploidy and 2 aneuploidies that were acquired *in vivo*. Chrs are shown from Chr1, Chr2-7, and R. **C.** Whole Chr LOH more frequently occurs on larger Chrs 1-3, and R in Gal^-^ isolates. Combinations of whole Chr LOH were not observed in Gal^+^ isolates.

**Table 2.** Summary of missegregation events by Chr.

| Chromosome | Whole chromosome aneuploidy |  |  |  | Whole Chr LOH |  |  |
| --- | --- | --- | --- | --- | --- | --- | --- |
|  | Total | 1N | 3N | 4N | Total | AA | BB |
| Chr1 | 18 | 10 | 7 | 1 | 81 | 81 | 0 |
| Chr2 | 16 | 10 | 6 | 0 | 28 | 18 | 10 |
| Chr3 | 11 | 10 | 1 | 0 | 41 | 11 | 30 |
| Chr4 | 24 | 9 | 12 | 3 | 8 | 8 | 0 |
| Chr5 | 34 | 10 | 24 | 0 | 6 | 5 | 1 |
| Chr6 | 83 | 5 | 72 | 6 | 4 | 4 | 0 |
| Chr7 | 19 | 5 | 12 | 2 | 5 | 5 | 0 |
| ChrR | 23 | 10 | 13 | 0 | 22 | 15 | 7 |

Number (events) for each genotype are shown separately for whole Chr aneuploidy (1N, monosomic, 3N, trisomic, 4N, tetrasomic) and whole Chr LOH (AA, homozygous allele A, BB, homozygous allele B)

Haploids were detected using flow cytometry optimized after detection of an initial haploid isolate from *in vitro* studies (HICKMAN *et al.* 2013). The detection of multiple haploid isolates (10/950 2DOG^R^ isolates recovered initially from the mouse oral cavity) was unexpected and exciting. Of note, only three haploids were perfectly euploid, with 7 being near-haploid; all the haploids tested were relatively unstable and readily converted to the autodiploid state (HICKMAN *et al.* 2015), and data not shown), suggesting additional haploids may have been present *in vivo*. We identified nine distinct haploid or near-haploid genotypes that were recovered from 6 different mice: 3 single haploids from 3 different mice (Fig.3B), 2 unique haploids from one mouse (D3M2), 3 distinct haploids from one host (D5M5), and 2 identical haploids (different CPs but treated as likely clones, D3M1) from one mouse (Table S2). Interestingly, only 3 genotypes were identical between Hickman *et al.* (HICKMAN *et al.* 2013) and this study, which suggests that the original isolates were a mixed population (supported by mixed flow cytometry profiles, data not shown) although the instability of haploids may have contributed as well.

Whole Chr LOH was detected for all Chrs, with higher frequencies seen for the larger Chrs (ChrR, 1-3) (Fig.4C, Fig.S4) (Table 2, S3). The frequency of missegregation events is not entirely a function of Chr size, however, as the frequency of events on Chr3 (1.8 Mb), was higher than either Chrs 2 (2.1 Mb) or ChrR (2.3 Mb), which are 2.1 and 2.3 Mb, respectively. Of note, whole Chr LOH of Chrs 2, 4, 6, and 7, was biased towards allele A, consistent with the failure to detect homozygous allele B in haploids or other isolates (FORCHE *et al.* 2008; HICKMAN *et al.* 2013; FORD *et al.* 2015; HICKMAN *et al.* 2015; HIRAKAWA *et al.* 2015; HIRAKAWA *et al.* 2017). This suggests that the B alleles of these Chrs harbor lethal recessive alleles (FERI *et al.* 2016) and therefore cannot be entirely lost. We did identify isolates with segmental LOH towards allele B for Chrs2, 4, and 6 (FIG.5A).

**FIG.5.**
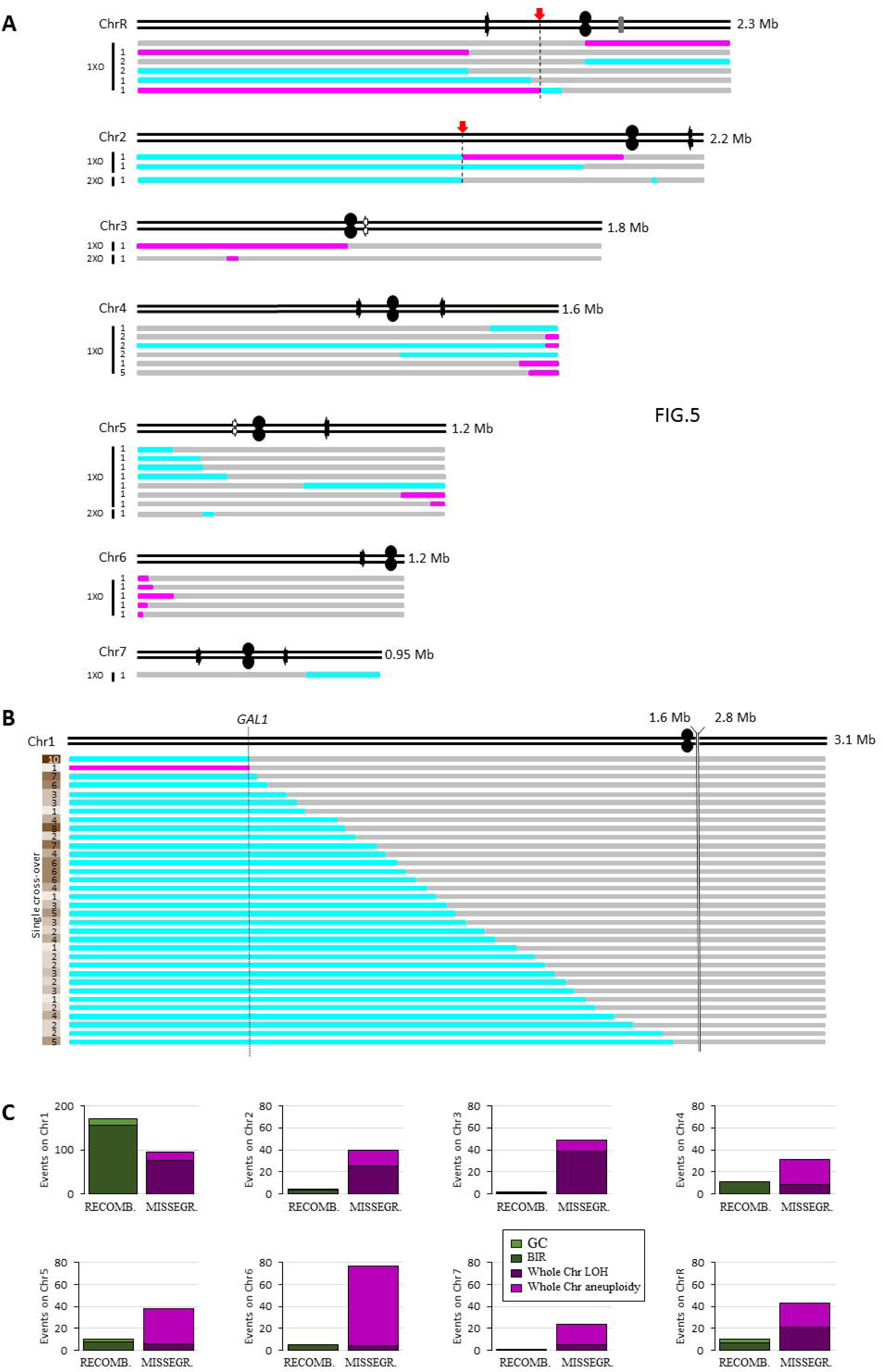
Recombination and missegregation events. **A.** Crossover-associated events most often lead to *GAL1* loss *in vivo*. **B.** Location of LOH breakpoints along Chr1. **C.** LOH breakpoints for Chr2-7, and R. Top horizontal black lines represent the two homologs; black oval represents centromeres, arrows show location of the major repeat sequence; Chr sizes are shown to the right of each Chr. The number of isolates for each genotype are indicated at the left; for Chr1 the numbers are in shades of yellow/brown, with higher numbers shaded darker; cyan, homozygous AA; magenta, homozygous BB; gray, heterozygous AB. Breakpoints were mapped in 25 kb bins. Exact start/end coordinates of break regions can be found in Table S4. Positions 1.6 - 2.8 Mb on Chr1 (indicated with 2 solid vertical black lines) are not shown due to lack of any LOH events across this region. XO, crossover, Maps are to scale.

## GAL1 LOH event characterization

The isolate collection illuminates the diversity of molecular mechanisms that can yield a Gal^-^ phenotype due to loss of the functional allele of *GAL1* from the B haplotype of Chr1 (Fig.5B). We classified 264 *GAL1* LOH events as either due to missegregation, (involving loss of the entire B Chr) or recombination (involving LOH across the subsection of Chr1L encompassing *GAL1*). Recombination was more frequent than missegregation (67% vs. 33% of the total events respectively) consistent with aneuploidy of large Chrs as rare or deleterious (Fig.5C) (Table S3). Among the Chr1 missegregation events, only the ten A-haplotype monosomies can be explained by a single step process: non-disjunction of the B copy of Chr1 during mitosis, leading to progeny with a single copy of the A homolog. The remaining aneuploids underwent at least two molecular events, either an increase in the copy number of the A version of Chr1 by missegregation, followed by loss of the B homolog, or vice-versa. Missegregation events were much more frequent than recombination on all other Chrs, with whole Chr aneuploidy more frequent on smaller Chrs (Chr5-7) and whole Chr LOH more prevalent on larger Chrs (Chr1-4, R) (Fig.5C).

Recombination events were categorized as: 1) LOH covering a Chr arm from the recombination initiation site through the telomere (likely break induced replication, BIR); 2) shorter-range LOH resulting from two crossover events that do not reach the telomere (double crossovers or gene conversions, GC); and 3) segmental copy number variations (CNVs) which includes truncations, amplifications and deletions. Recombination events on Chr1L were dominated by BIR (94%), with GC events implicated in the remainder (6%, Fig.5B and 6). Rare recombination events also were detected on all other Chrs in a few Gal^+^ as well as in Gal^-^isolates and BIR was far more frequent than GC (Fig.5A and 6).

**FIG.6.**
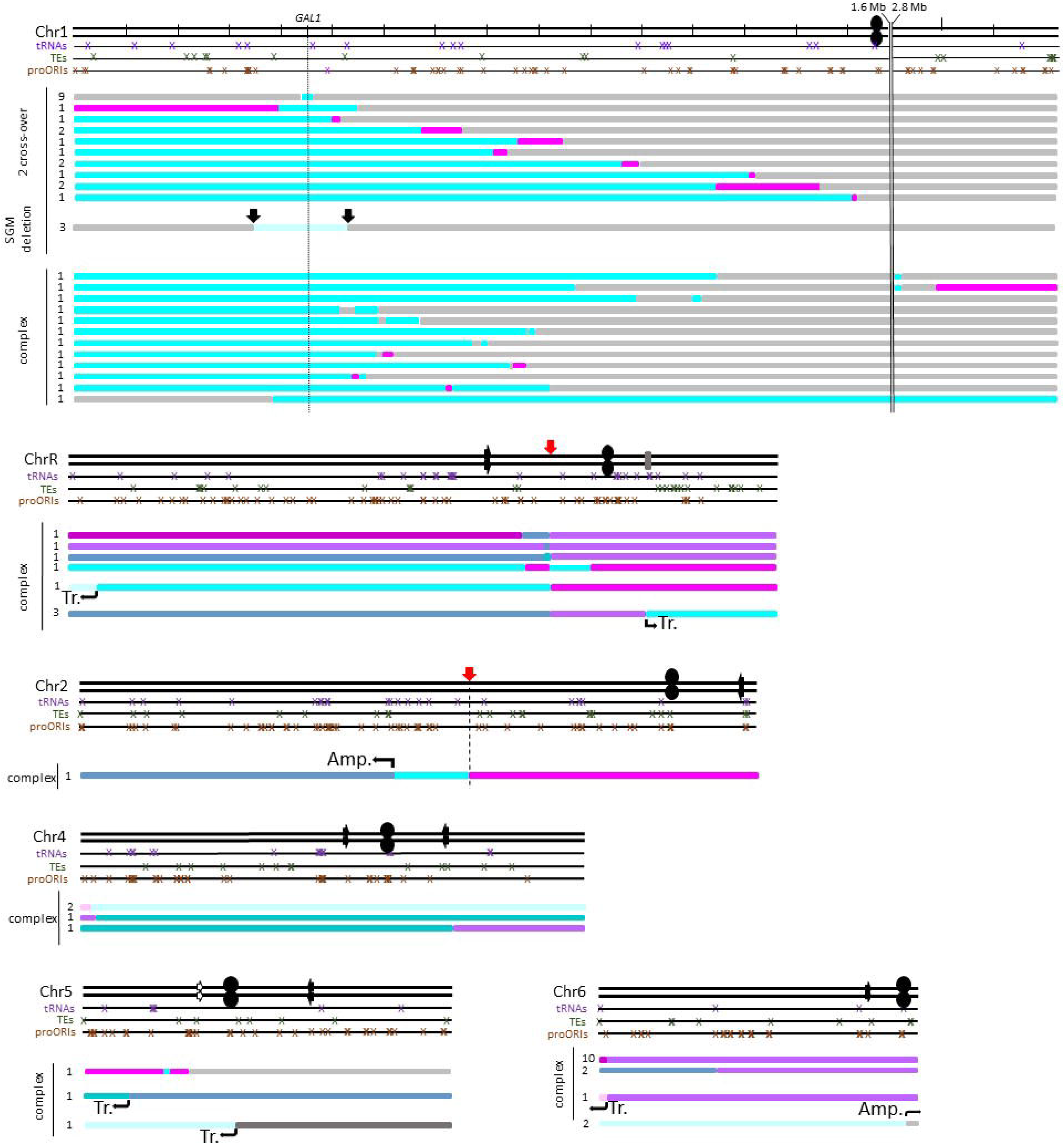
Complex changes on individual Chrs include multiple recombination events on single Chrs (mostly Chr1), segmental deletions, truncations, and amplifications. For legend, please see Fig.5.

The sites of recombination across Chr1L appeared relatively randomly distributed between *GAL1* and *CEN1* (Fig.5B). Four potential hot spot regions were identified by binning Chr1L breakpoints every 50 kb along Chr1L (FIG.S5): two to the right of the *GAL1* (at ∼450 kb) locus (451-500 kb and 501-550 kb), one between 701-750 kb and the last one between 851-900 kb (Fig.S5, Table S4). Most of these break regions were not near any genome elements known to promote double strand breaks such as transposable elements.

In addition to the missegregation and recombination patterns described above, a small number of complex rearrangements were also detected. These included Chr truncations and recombination events that involved segmental aneuploidies and could only arise through multiple sequential events on a single Chr (Fig.6). Complex events were only seen following 2DOG selection and were most frequent on Chr1 (Fig.6) (see Table S4 for break coordinates). In addition, several isolates had multiple crossover events on Chr1, some of them involving both Chr arms with LOH to AA and BB alleles at different positions (Fig.6). Taken together, these complex genotypes suggest that double-strand breaks (DSBs) were repaired via multiple, distinct mechanisms (see discussion).

## Recurrent events

The finding that the same genomic changes appeared in multiple mice supports the idea that these are general responses of the genome to conditions encountered in the oral cavity during early stages of infection. Many missegregation events occurred in multiple mice (Fig.7, S6, Table S5). Homozygosis of Chr1 to the AA genotype was seen in every mouse following 2DOG selection, presumably because this is an efficient mechanism for *GAL1* LOH (Fig.4). Other recurrent whole Chr events, such as trisomy of Chr6, which appeared in the Gal^+^ isolates as well (Fig.S6), was unexpected. Uniparental disomy of Chr3 and Chr5 trisomy were also prevalent. Whether these different missegregation events are advantageous during early infection or during the transition into and out of the host, remains to be determined.

**FIG.7.**
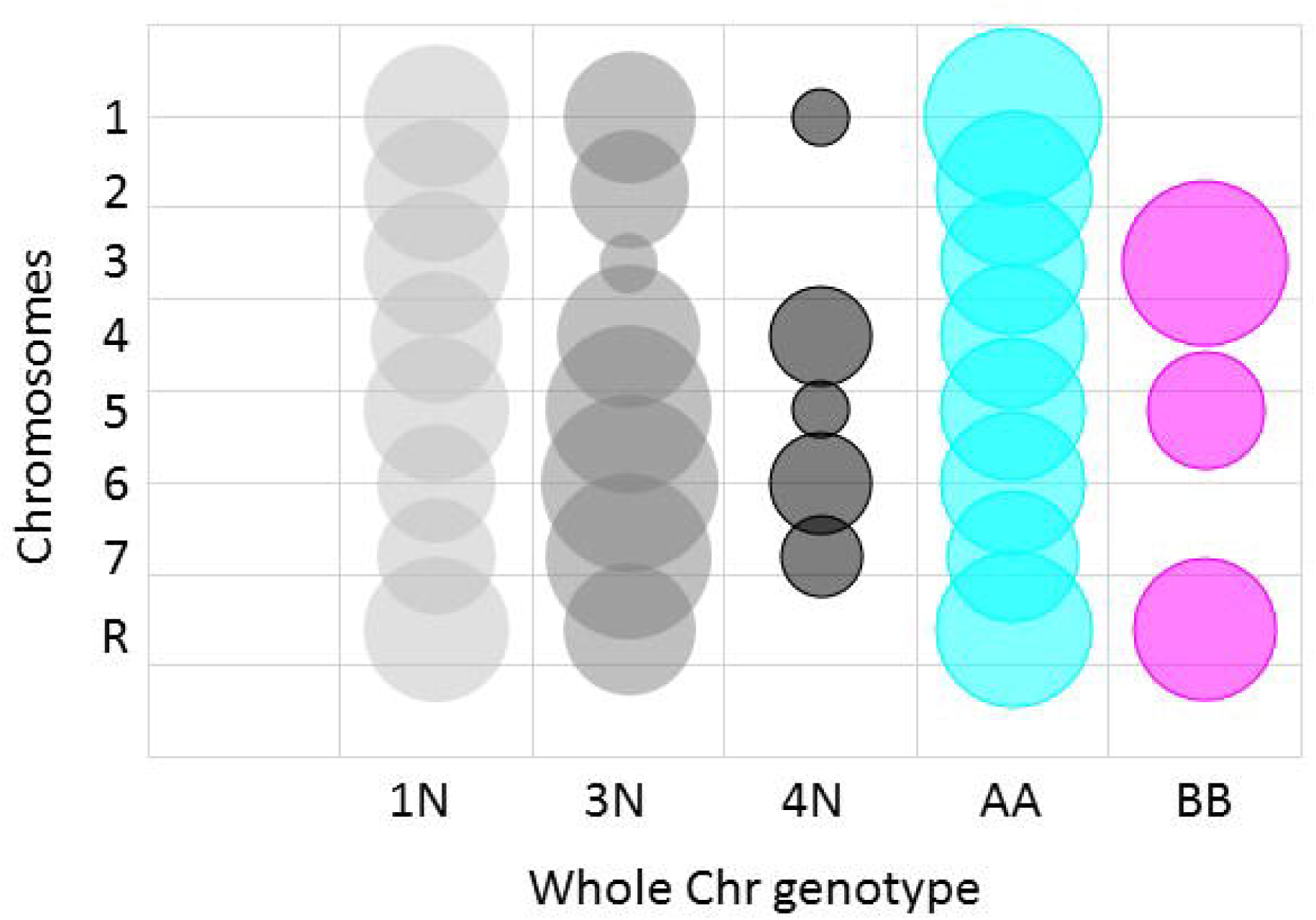
Recurrent missegregation events are frequent. Calculations were done for mice with *C. albicans* populations size ≥ 12 (9 of 17 mice); bubble sizes reflect the percent mice where the specific missegregation event (indicated on x-axis) was found. For example, whole Chr1 LOH allele AA and whole Chr6 trisomy were found in all 9 mice (100%). Y-axis, Chr1-7, and R; xaxis, missegregation genotypes.

## Hypervariability in evolved isolates

Multiple combinatorial (i.e, recombination + missegregation) events were most frequent in Gal^-^ isolates that also exhibit CPs (Fig.8A). Not surprisingly, multiple missegregation events were found together much more frequently than multiple recombination events—likely because aneuploidies and LOH arise during concerted Chr loss from tetraploid intermediates (FORCHE *et al.* 2008; HICKMAN *et al.* 2015). Recombination events were less prevalent in general and correspondingly the frequency of multiple recombination events was also much smaller.

**FIG.8.**
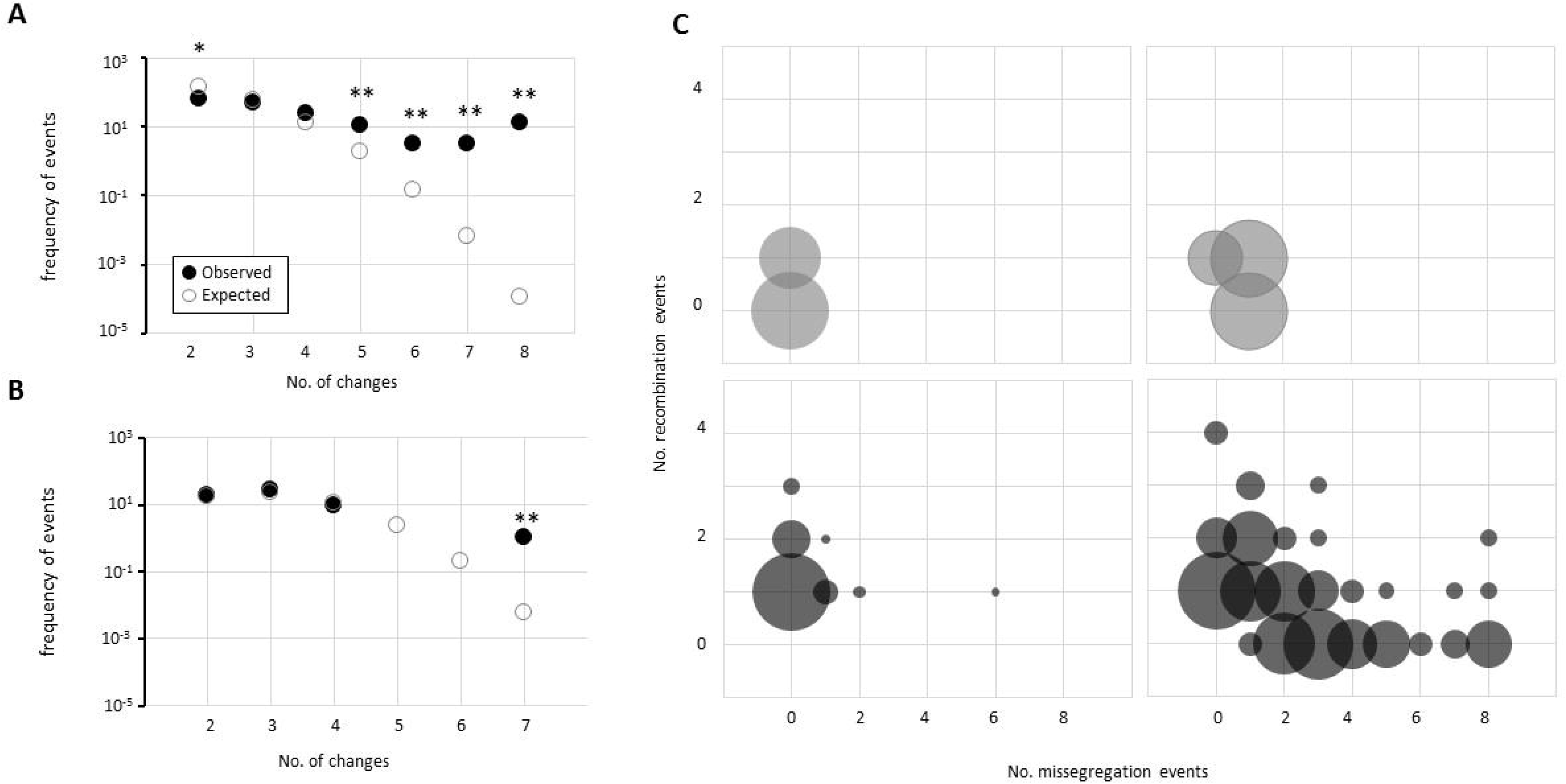
Multiple changes (> 5) per isolate are significantly more frequent than what would be expected by random chance alone *in vivo* but not *in vitro* **A.** Multiple combinatorial (recombination (REC) + missegregation (MIS)) events are most frequent in Gal^-^ with CPs. Percent of multiple event types for Gal^+^ isolates (top left), Gal^+^ plus CP (top right), Gal^-^ (bottom left) and Gal^-^ plus CP (bottom right). Y-axis, number of recombination events/isolate; x-axis, number of missegregation events per isolate. Bubble size represents the number of isolates with indicated combinations, e.g. number of isolates that have 1 recombination and 1 missegregation event. Expected versus observed frequencies of changes *in vivo* (**B.**) and *in vitro* (**C.**). Significance is indicated by ** (p = 0.01).

We previously found that tetraploid isolates that underwent Chr loss yielded progeny with evidence of mitotic recombination that tended to involve multiple events on different Chrs (FORCHE *et al.* 2008). Combined with our observations of multiple and/or complex changes per isolate (Fig.5A and B, Fig.6), this suggests that once an isolate has undergone one mutational change it has an increased likelihood of additional changes. To quantitatively explore this idea, we calculated the frequencies of multiple *vs.* single events detected in the *in vivo* and *in vitro* samples (data not shown), as well as the frequency expected if each event arose randomly. The frequency of events was significantly greater than random for genome change *in vivo* but not *in vitro* (only ≥ 7 changes was significant) (FIG.8B and C, Table S4), indicating that in the isolates studied, highly diverse isolates are overrepresented. This implies that rare individuals undergo high levels of recombination that involve multiple Chrs.

## Discussion

To understand the evolutionary forces responsible for genomic rearrangements leading to fitter genotypes, one must first identify the types of changes that reshape the genome (CHADHA AND SHARMA 2014). Here, we provide the first population-level study of the standing variation that arises in *C. albicans* during oropharyngeal candidiasis by analyzing several hundred isolates recovered from 17 mice at different time points during the infection. Importantly, this study design provided the perspective of time within the host. Flow cytometry and ddRADseq of 429 isolates detected many types of events due to missegregation, recombination and multiple events of both types. Our observations of missegregation and DSB-associated changes are consistent with two recent studies of genotypic and phenotypic intra-species variation and the evolution of drug resistance in single isolates of clinical *C. albicans* isolates (FORD *et al.* 2015; HIRAKAWA *et al.* 2015). While previous studies provide an important snapshot of ongoing changes in human infections, the lack of multiple isolates per time point makes it very difficult to recapitulate isolate genealogies throughout evolution. Of note, diversity was detectable even one day post infection, suggesting that either changes arise rapidly upon the shift in growth conditions from liquid medium to the mouse and back (JACOBSEN *et al.* 2008) or that exposure to the host environment for only 24 h of infection is sufficient to induce genotypic changes.

The detection of multiple independent haploid or near-haploid isolates with different genotypes was surprising, suggesting that haploidization repeatedly occurs in the oral cavity. We previously found Chr missegregation in isolates recovered after passage in a systemic model of infection and after *in vitro* exposure to physiologically relevant stressors (FORCHE *et al.* 2011). *In vitro,* the length of LOH tracts (short, long, whole Chr) was associated with the type and severity of stress applied. Here, all three types of LOH arose at appreciable frequencies along with high levels of aneuploidy, supporting the idea that *C. albicans* is exposed to significant combinatorial stress in the oral cavity even though it appears to flourish in the oral cavity during oropharyngeal candidiasis.

We detected a positive correlation between specific CPs and Chr missegregation. A large proportion of CPs were small in diameter and had completely smooth or less wrinkly colonies (Fig.2C), suggesting that they grow less well than the parental strain under the conditions tested and have defects in filamentous growth. This is reminiscent of the slow growth seen for aneuploidy *Saccharomyces cerevisiae* isolates grown in lab media (TORRES *et al.* 2007; THORBURN *et al.* 2013), which is thought to be the result of unbalanced protein stoichiometry, difficulty segregating aneuploidy Chrs or higher demands for DNA replication (STORCHOVA *et al.* 2006; TORRES *et al.* 2008; PAVELKA *et al.* 2010; TORRES *et al.* 2010; BENNETT *et al.* 2014; HIRAKAWA *et al.* 2015).

Mutants with filamentation defects cause less damage to epithelial and endothelial cells *in vitro* (PHAN *et al.* 2000; TSUCHIMORI *et al.* 2000; BENSEN *et al.* 2002). This suggests that the isolates with reduced filamentous growth may not express hyphal-specific genes (e.g., *ALS3*, *SAP4* and *SAP6*) and/or may not be recognized as readily by the host immune cells. The majority of isolates with small CM acquired whole Chr aneuploidy, supporting the idea that these isolates may grow slowly under standard lab conditions, yet might have an advantage *in vivo* (SEM *et al.* 2016). Interestingly, a subset of isolates recovered after a systemic infection in mice also exhibited aneuploidy and LOH (FORCHE *et al.* 2009a).

Chr6 trisomy was much more frequent than other aneuploidies, and Chr6ABB was twice as frequent as Chr6AAB. Chr6 harbors multiple members of important virulence gene families, such as secreted aspartic proteases, lipases and adhesins (HUBE *et al.* 2000; NAGLIK *et al.* 2004; SCHALLER *et al.* 2005; HOYER *et al.* 2008; DJORDJEVIC 2010), the *NAG* gene cluster important for alternative carbon utilization (KUMAR *et al.* 2000) and *RAD52*, a gene important for DSB repair (CIUDAD *et al.* 2004; CIUDAD *et al.* 2005). Interestingly, overexpression of Rad52 increased genome instability (TAKAGI *et al.* 2008). Therefore, an extra copy of *RAD52* could potentially lead to increased genome instability and amplification of specific advantageous alleles (e.g. one extra copy of allele A) could promote adaption to specific environments such as the oral cavity. Follow-up experiments will test the effect of Chr6 trisomy on survival, persistence, and virulence of *C. albicans* in the oral cavity.

DSBs arise from endogenous sources including reactive oxygen species (e.g., produced by immune cells), collapsed replication forks, and from exogenous sources including chemicals that directly or indirectly damage DNA (SHRIVASTAV *et al.* 2007). The utilization of the *GAL1* selection system not only allowed us to identify the major classes of genome changes and to catalogue the types of LOH events that resulted in a Gal^-^ phenotype, but it also enabled us to make hypotheses about the types of mechanisms that are involved in DSB repair. While the majority of LOH was likely the result of BIR with or without crossover, more complex LOH events also arose in a subset of isolates (see Fig.5C). The LOH signatures on Chr1 are consistent with what would be observed after short and long patch mismatch repair using different alleles as repair templates (COÏC *et al.* 2000; MARTINI *et al.* 2011; BOWEN *et al.* 2013). Furthermore, more than one mismatch machinery may have been involved in repairing breaks. Strikingly, similar LOH signatures were observed during mitotic DSB repair in *S. cerevisiae* (GUO *et al.* 2017; HUM AND JINKS-ROBERTSON 2017), suggesting that these mechanisms may have been conserved through evolution.

Whether genotypic variation arises through the parasexual cycle or via mitoic defects followed by Chr missegregation accompanied by recombination events remains an outstanding question. Our previous analysis of parasexual progeny showed that the majority of them were aneuploid, that Chr missegregation predominated, that changes were observed for multiple Chrs and that several isolates had short recombination tracts on multiple Chrs (FORCHE *et al.* 2008). A more recent study examined 32 parasexual progeny generated *in vitro* for a wide range of virulence-associated traits and showed that parasexual mating can generate phenotypic diversity *de novo*, and has important consequences for virulence and drug resistance (HIRAKAWA *et al.* 2017) Direct evidence for the parasexual cycle *in vivo*, however, remains elusive and the mechanism of mitotic failure followed by Chr loss events cannot be ruled out (HARRISON *et al.* 2014).

Importantly, here we identified a substantial level of highly variable isolates, higher than what one would expect by random chance alone. We hypothesize that hypervariable subpopulations may be present in many natural populations, and that this diversity can enable rapid adaptation in time of stress or environmental stochasticity. Whether the observed changes are beneficial, detrimental or neutral remains to be determined, and is likely to be specific to the particulars of the environment. The link between how specific genotypic changes affects survival, persistence, and the virulence potential of *C. albicans,* and whether the host recognizes and responds to this variation remains to be discovered.

## ACKNOWLEDGEMENTS

We thank Carter Meyers for assistance with the animal studies and Darren Abbey for the MAtlab script to determine likelihood for all permutations for each number of genotypic changes. AMD, GC, EJ and AF were funded by NIH grant R15 AI090633 to AF. SGF was funded by NIH grant R01DE022600 and JB was funded by an European Research Council Advanced Award 340087 (RAPLODAPT). ACG was supported by a CIHR Banting Postdoctoral Fellowship.

---

FIG.S1. Detailed experimental overview. This figure is an expansion of Fig.1 and includes detailed information about the total number of Gal^+^ (YPD, no selection) and Gal^-^ (2DOG, selection) isolates that were analyzed, that exhibit CPs, and genomic changes. Numbers are parsed by mouse and the time that isolates spent in the mouse (in days). In addition, this figure provides the actual total frequencies for CPs and genome changes determined by extrapolation from the total number of CFUs.

FIG.S2. The frequencies of *GAL1* LOH is 2 orders of magnitude higher *in vivo*. Each open circle represents one mouse (*in vivo*) or independent *in vitro* cultures (grown for 16 hrs).

FIG.S3. Single and double aneuploidies are detected for both Gal^+^ (YPD, no selection) and Gal^-^(2DOG, selection) isolates. Shown are examples for isolates with 1 aneuploidy (1AN) and 2 aneuploidies (2AN) that were acquired *in vivo*. Chrs are shown from Chr1, Chr2-7, and R, gray and black colors were used to show start and end of each Chr.

FIG.S4. Whole Chr LOH is more frequent for larger Chr1-3, and R. Shown are examples for whole Chr LOH either toward allele A or allele B and the number of isolates that acquired these specific genome changes. * For Chrs4 and 7 no isolate with a single whole Chr LOH was observed. Chrs exhibiting whole Chr LOH are boxed.

FIG.S5. Mapping of LOH breaks along Chr1L reveals 4 hotspot regions. Each of these regions contains between 15 and 22 breaks and is marked with a red arrow. Breaks were mapped in 25 kb bins due to low resolution of ddRADseq. *CEN1*, centromere 1

FIG.S6. Missegregation events are highly recurrent across mice. Shown are summaries for 9 mice with N > 12. Chrs are indicated on the y-axis. Mouse IDs are indicated across the top. For each mouse there are two columns of pie charts. The first column shows the number of Chrs that are 1N, 2N, 3N, and 4N with shades of brown going from light (1N) to dark (4N). The right column shows allele status (heterozygous (gray), allele A (cyan), allele B (magenta)).

